# Epigenetic Potential and DNA Methylation in an Ongoing House Sparrow (*Passer domesticus*) Range Expansion

**DOI:** 10.1101/2020.03.07.981886

**Authors:** Haley E. Hanson, Chengqi Wang, Aaron W. Schrey, Andrea L. Liebl, Rays H. Y. Jiang, Lynn B. Martin

**Author notes:** **Corresponding Author:** To whom correspondence may be addressed. University of South Florida, Global and Planetary Health, Tampa, FL. or. **Author Contributions:** H.E.H., A.W.S., and L.B.M. designed research. A.L.L. collected samples. H.E.H. and A.W.S. performed research. C.W. analyzed data. H.E.H. wrote the first draft. All authors contributed to manuscript revisions.

## Abstract

During range expansions, epigenetic mechanisms may mediate phenotypic responses to environmental cues, enabling organisms to adjust to novel conditions at novel sites. Here, we predicted that the number of CpG sites within the genome, one form of epigenetic potential, would be important for success during range expansions because DNA methylation can modulate gene expression and hence facilitate adaptive plasticity. Previously, we found that this same form of epigenetic potential was higher in introduced compared to native populations of house sparrows (*Passer domesticus*) for two immune genes (Toll-like receptors *2A* and *4*). Here, we took a reduced-representation sequencing approach (ddRadSeq and EpiRadSeq) to investigate how CpG site number, as well as resultant DNA methylation, varied across five sites in the ∼70 year-old Kenyan house sparrow range expansion. We found that the number of CpG sites increased towards the vanguard of the invasion, even when accounting for variation in genetic diversity among sites. This pattern was driven by more losses of CpG sites towards the core of the invasion (the initial site of introduction). Across all sequenced loci, DNA methylation decreased but became more variable towards the range-edge. However, in the subset CpG sites proximal to mutated CpG sites, DNA methylation increased and variation declined. These results indicate that epigenetic potential influenced the Kenyan house sparrow range expansion, likely by providing greater phenotypic plasticity which is genetically assimilated as populations adapt to local conditions. Similar mechanisms might underlie the successes and failures of other natural and anthropogenic range expansions.

## Introduction

Epigenetic modifications, such as DNA methylation, play a critical role in linking environmental variation to phenotypic variation by modifying how genes are expressed (1). In vertebrates, these processes are instrumental to cellular and tissue differentiation during development, but these same modifications can also affect evolutionarily-relevant behavioral, morphological, and physiological plasticity (2, 3). As epigenetic variation, including DNA methylation patterns, predominates in particular genomic regions (e.g., CpG dinucleotides), genomes might differ in their capacity to be modified epigenetically (4). When genomic variation associated with epigenetic marks occurs in genes that affect fitness, natural (and neutral) selection should follow, leading to differences in epigenetic potential among individuals, populations, and species (4). Consequently, epigenetic potential (i.e., the capacity for epigenetic mechanisms to capacitate phenotypic variation) might become common or rare depending on ecological and evolutionary context and the value of phenotypic plasticity in a given area (5). For instance, during range expansions, individuals face relatively novel conditions that require rapid phenotypic responses, but also risk reduced performance because of low population genetic diversity or founder effects (6). In such situations, we expect epigenetic potential to predominate. As generations reproduce and experience selection in colonized environments, though, the local value of epigenetic potential might wane as mutations occur, selection ensues, and phenotypically plastic genotypes are outcompeted by genotypes with genetically canalized adaptations (5). In this light, we asked if epigenetic potential contributed to a successful range expansion of one of the most ubiquitous avian species, the house sparrow (*Passer domesticus*) (Fig. 1).

**Figure 1:**
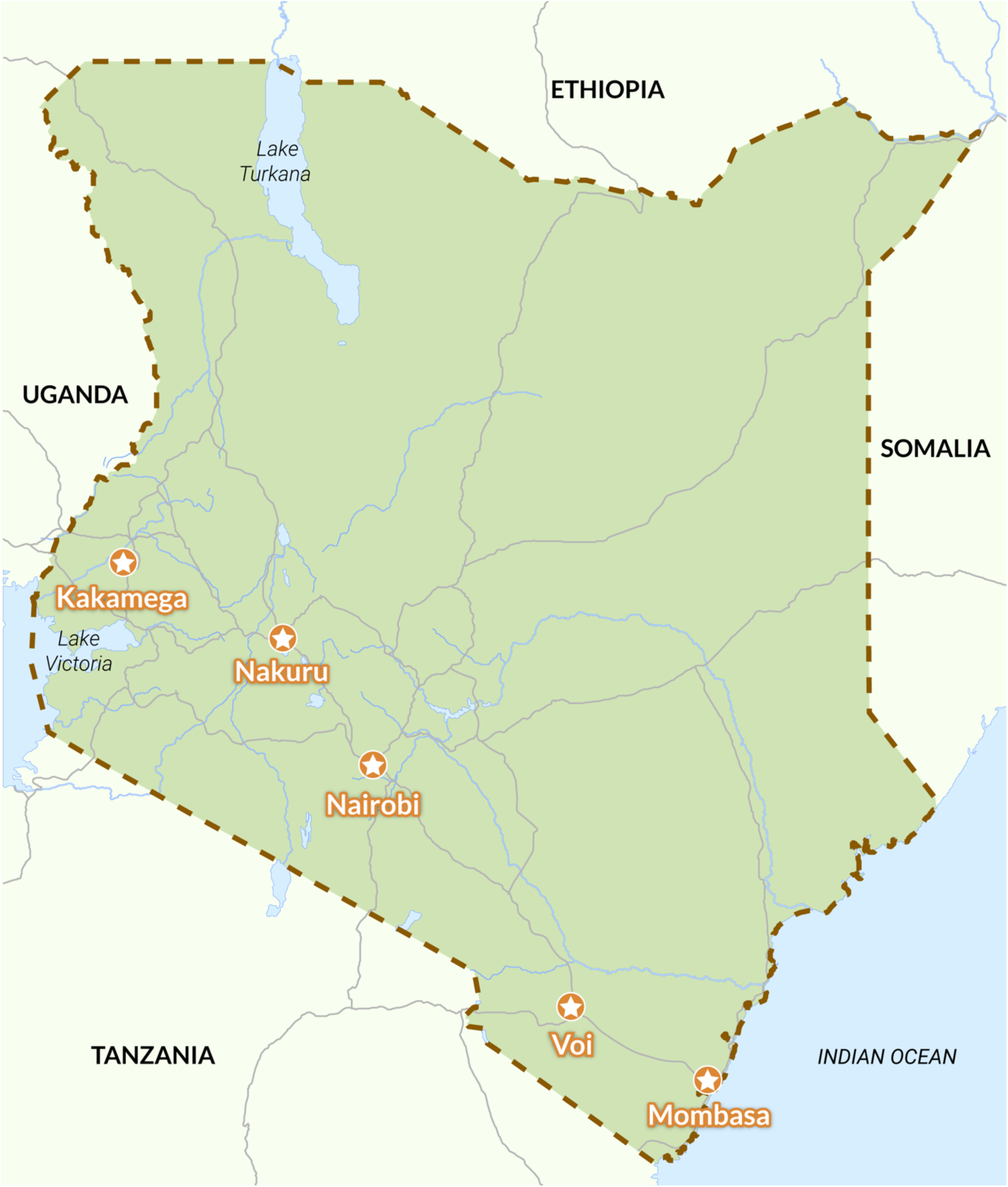
Map of cities sampled in Kenya.

House sparrows (*Passer domesticus*) moved out of the Middle East thousands of years ago as humans colonized Europe and over the last 170 years have achieved a near global distribution largely due to intentional or accidental movements by humans (7, 8). As house sparrows spread globally, they have had to cope with a wide range of biotic and abiotic factors. Concurrently, some introduced groups would have faced genetic challenges associated with introduction events and expansions, namely founder effects and bottlenecks (7). Despite these challenges, house sparrows endure and exhibit extensive phenotypic variation across much of the globe (9–11). In one of their most recent range expansions, Kenya, house sparrow trait variation, including the regulation of glucocorticoid hormones and immune genes and several behaviors associated with invasions, tracks with relative population age (12–17). This paradox of extensive trait variation when genetic variation is comparatively low (relative to native populations) might be resolved by epigenetic compensation, which we observed previously in this system (18). In other words, when population genetic diversity was low at a capture site along the invasion, epigenetic diversity was high (18, 19).

Epigenetic potential may manifest through multiple mechanisms, but here we focus on the number of genetic motifs upon which DNA can be methylated (5, 20, 21). DNA methylation occurs when a methyl group is added to the fifth carbon position of a cytosine within a cytosine-phosphate-guanine site, or CpG site (1). DNA methylation can be induced or eliminated rapidly following stimulation from a range of factors (e.g., diet, transcription factor activity, stress, etc.), and it can induce, inhibit, or enhance gene expression depending on where it occurs in the genome (1, 20, 22). In principle, each CpG site represents an opportunity for DNA methylation to alter gene expression (21). Previously, we found that DNA methylation affected the expression of one immune gene, Toll-like receptor 4 (*TLR4*), a gene that also varied in expression across the Kenyan range, being expressed most at the range edge (23, 24). Here, we asked whether epigenetic potential varied across this same range expansion, investigating specifically how CpG number and DNA methylation state varied among sites of different age. This work was partly motivated by our recent discovery that the number of CpG sites for two immune genes (Toll-like receptors *2A* and *4*) was higher in introduced compared to native house sparrows (23). There, we argued that epigenetic potential was higher for these two genes in introduced populations because it would allow for more plasticity in immune responses, an important trait in areas where pathogens would have had a short evolutionary history with sparrows.

In the present study, we used reduced representation library-based sequencing to compare CpG number and DNA methylation among five geographic locations spanning the ∼70 year old Kenyan range expansion (25). Genetically, we queried i) how the number of CpG sites changed across the range expansion; and ii) whether CpG sites were being lost or gained and if so, whether CpG mutations were more common than other mutations. Epigenetically, we asked iii) whether global DNA methylation levels varied amongst sampling sites; and iv) whether CpG site losses and gains were related to nearby methylation. We predicted that epigenetic potential would increase towards the range edge with range edge sparrows gaining more or losing fewer CpG sites than those captured closer to the site of introduction (range core). We also expected more epigenetic potential to underlie more methylation, yet the complex interplay among CpG sites and DNA methylation status made more precise predictions along the range expansion impossible.

## Results

House sparrow hippocampal samples were collected from birds captured in five cities across southern Kenya. House sparrows were initially introduced to the port city of Mombasa, and, as in previous studies, we used distance from Mombasa as a proxy for time since colonization (see 9, 10, 19, 20). Birds were captured from Mombasa (0 km), Voi (160 km), Nairobi (500 km), Nakuru (650 km), and Kakamega (885 km). Screening sparrows from these five cities, we conducted double digest Restriction-site Associated DNA sequencing (ddRAD-seq) with a traditional restriction enzyme (*MspI*) and a methylation-sensitive restriction enzyme *(HpaII*-EpiRADseq) (n= 64, sample sizes listed in *SI Appendix*, Table S1). *MspI* and *HpaII* cut DNA at the same sequence (CCGG), but differ in their sensitivity to methylation, allowing us to calculate population genetic statistics and compare the presence or absence of methylation at loci among individuals (25). Following quality control, ddRADseq returned 452,465 reads and 1,205 single nucleotide polymorphisms (SNPs). We chose to sample hippocampus in the hopes of capturing DNA methylation data within genes relevant to range expansions, such as those related to memory and behaviors relevant to exploration of novel environments, but unfortunately the coarse nature of this sequencing approach and the lack of an annotated house sparrow genome made it so very few specific genes or gene regions could be identified (see Materials and Methods) (5).

### CpG Sites Across the Range Expansion

CpG sites were relativized using GpC sites for total, losses, and gains of CpG sites, as well as for mutation types (see Materials and Methods for equations) (28). The number of relativized CpG sites increased with distance from Mombasa (Fig. 2*A*; R= 0.49, F-test, p < 0.0001). We identified 118 SNPs causing a CpG site to be gained across 28 unique genomic locations, and 191 SNPs causing a CpG site to be lost across 36 unique genomic locations. We detected a significant difference in the number of CpG sites being lost across the range expansion with more CpG sites lost towards the range core (Fig. 2*B*; R= −0.51, F-test p < 0.0001). There was no difference in the number of gains among sites (*SI Appendix*, Fig. S1; R= −0.1, F-test p= 0.41). In regard to the losses of CpG sites, we found that CpG to CpA and TpG mutations were the most frequent types of mutations overall (*SI Appendix*, Fig. S2 and Table S2; CpA FDR <0.05, TpG FDR <0.1). No other mutation type was significantly different from any other form (*SI Appendix*, Fig. S2 and Table S2). Along the range expansion, CpA and TpG mutations increased towards the range core, opposite of the pattern we observed for CpG number (Fig. 2*C* and *D*; CpA: R= −0.40, F-test p < 0.001; TpG: R= −0.40, F-test p = 0.001).

**Figure 2:**
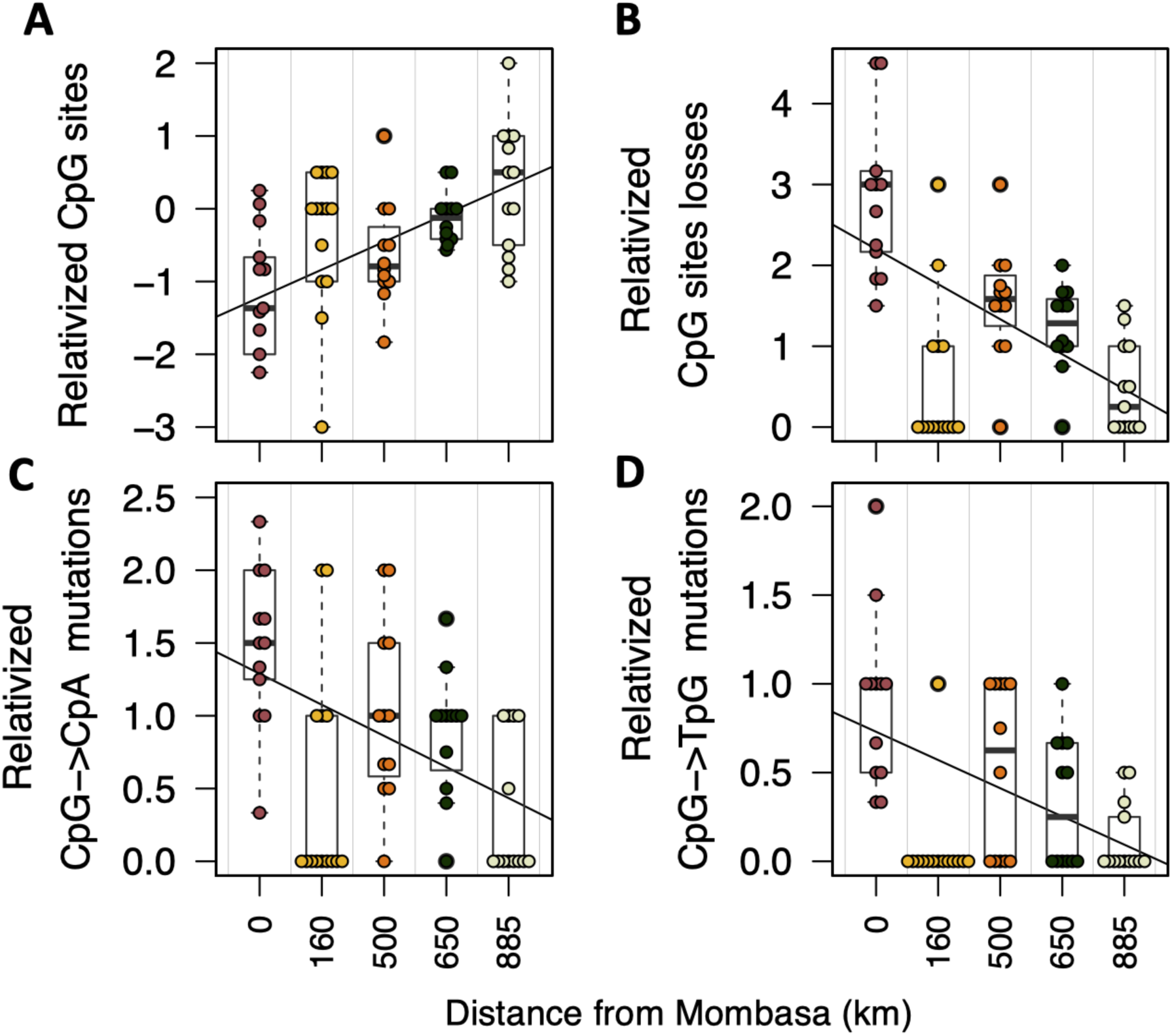
CpG sites, losses of CpG sites, and prevalence of CpA and TpG mutations across the Kenyan house sparrow range expansion. (A) CpG sites increase towards the range edge (R= 0.49, F-test, p < 0.0001). (B): Losses of CpG sites decrease towards the range edge (R= −0.51, F-test p < 0.0001). CpA (C) and TpG (D) mutations decrease towards the range edge (CpA: R= −0.40, F-test p < 0.001; TpG: R= −0.40, F-test p = 0.001). In A-D, points represent individual samples (n= 64). CpG sites were relativized using equations found in *Materials and Methods*.

### Methylation Across the Range Expansion

Out of 14,659 loci, 4,518 loci exhibited differential methylation between pairs of sampling sites. Across all sequenced loci, DNA methylation declined towards the range edge (Fig. 3*A*; R= −0.17, F-test p = p < 0.0001), but variation in methylation across loci increased towards the range edge (*SI Appendix*, Fig. S3*A*; R= 0.06, F test p < 0.0001). Of the 4,518 loci that exhibited differences in DNA methylation, 58 were near a CpG site that had been lost or gained in at least one bird (i.e., within the same read segment, median length of 310 base pairs). DNA methylation in this subset of CpG sites increased towards the range edge (Fig. 3*B*; R= 0.20, F test p= 0.025), but variation in DNA methylation status, however, decreased (*SI Appendix*, Fig. S3*B*; R= −0.21, F test p= 0.003).

**Figure 3:**
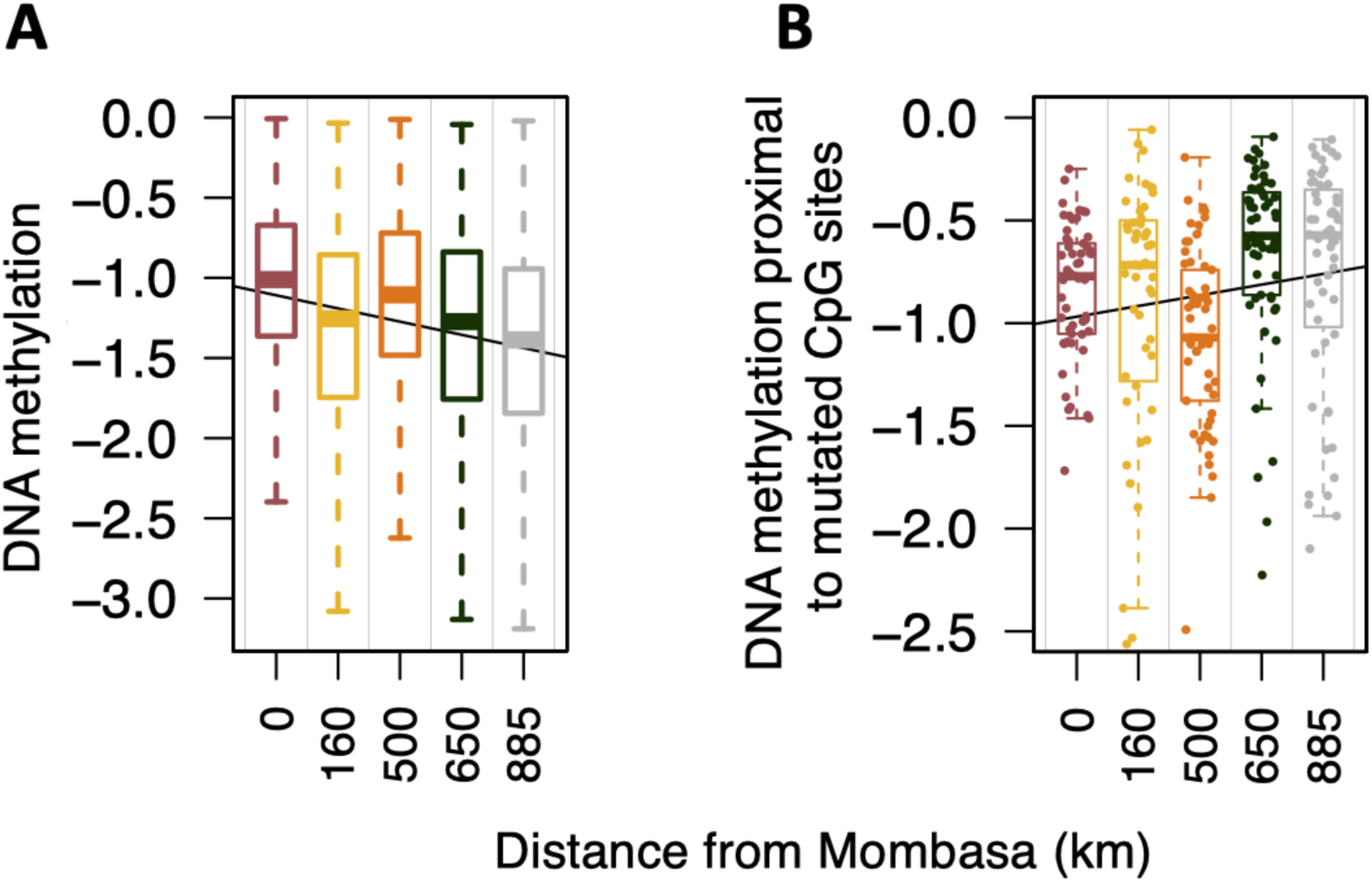
DNA methylation across the Kenyan house sparrow range expansion. (A) DNA methylation measured across all sequenced loci decreased towards the range edge (R= −0.17, F-test p = p < 0.0001). (B) DNA methylation at loci proximal to a CpG site mutation increased towards the range edge (R= 0.20, F test p= 0.025). In (B), points represent loci and are plotted with horizontal “jitter” (n=58). Points were not reported in (A) as n= 4,518.

### Genetic Diversity and Population Structure

Among the five Kenyan sites, observed heterozygosity ranged from 0.160 to 0.258, expected heterozygosity ranged from 0.125 to 0.187, number of private alleles ranged from 0 to 31, and the inbreeding coefficient ranged from −0.115 to −0.029 (Table 1). Neither observed heterozygosity, number of private allelic sites, nor the inbreeding coefficient were related to distance from Mombasa (*SI Appendix*, Fig. S4). A Discriminant Analysis of Principle Components (DAPC) predicted only two genetic clusters, yet both clusters contained at least one individual from every city. Thus, there was no evidence of genetic constraints (e.g., genetic diversity) or artifacts (e.g., founder effects) underlying the above geographic patterns in CpG number or methylation (*SI Appendix*, Fig. S5).

**Table 1:** House sparrow sampling locations across Kenya and estimates of observed heterozygosity (H_o_), expected heterozygosity (H_e_), private allelic sites, and the inbreeding coefficient (F_is_).

| City | Distance from Mombasa (km) | Observed Heterozygosity ($H_o$) | Expected Heterozygosity ($H_e$) | Private Sites | Inbreeding Coefficient ($F_{is}$) |
| --- | --- | --- | --- | --- | --- |
| Mombasa | 0 | 0.258 | 0.187 | 31 | -0.105 |
| Voi | 160 | 0.196 | 0.136 | 0 | -0.115 |
| Nairobi | 500 | 0.160 | 0.125 | 13 | -0.029 |
| Nakuru | 650 | 0.193 | 0.133 | 19 | -0.108 |
| Kakamega | 885 | 0.198 | 0.148 | 11 | -0.072 |

## Discussion

We found that house sparrows at the vanguard of a range expansion maintained more CpG sites (relativized CpG sites) than those near the core of the invasion. Birds near the range edge did not gain CpG sites, rather individuals towards the initial site of introduction lost more CpG sites. Further, CpG sites that were lost were associated with high levels of DNA methylation; also, CpG mutations (to CpA or TpG) were more common than any other mutation. Epigenetically, we found that DNA methylation across all sequenced loci decreased towards the range edge, but variation in DNA methylation increased towards the edge. By contrast, at CpG sites that were lost or gained via mutation, we observed more DNA methylation and less variation among individuals towards the range edge (*SI Appendix*, Fig. S3*A*). Finally, there was no evidence that the above patterns in CpG numbers and resultant methylation were driven by underlying genetic patterns (6). Indeed, no population genetic measures were related to population age, nor was there evidence of extensive genetic differentiation among sites (Table 1, *SI Appendix*, Fig. S4 and S5). Below we discuss the ramifications for these results for house sparrow invasions as well as the role of epigenetic potential in other anthropogenic and natural range expansions broadly.

### Epigenetic Potential Underlying Phenotypic Plasticity

CpG sites increased towards the expanding edge of the Kenyan house sparrow invasion (Fig. 2*A*). House sparrows arrived in Kenya in the 1950s from an earlier introduction to South Africa that occurred at the turn of the 20^th^ century, which reduced genetic diversity substantially compared to populations elsewhere in the world (18). Previous work, relying on highly mutable microsatellites, revealed that Kenyan house sparrows within the southern half of the country were colonized from a single founding source (26, 29). Further, genetic diversity increased towards the range edge and correlated strongly with distance to Mombasa (26). Here, however, relying on a larger but less rapidly evolving subset of the genome, we detected no population genetic structure, nor relationships between population genetic traits (e.g., heterozygosity, inbreeding) and distance from Mombasa (Table 1, *SI Appendix*, Fig. S4). Although we cannot conclude that genetic diversity and founder effects explain none of the variation in epigenetic potential and methylation we observed, such a striking geographic pattern and the manner in which CpG prevalence was eroded via CpG-specific mutations suggest some support for our original hypothesis (Fig. 2*A*). In other words, we propose that this particular pattern represents evidence that epigenetic potential facilitates phenotypic plasticity, which is later genetically canalized to fix gene expression at particular levels.

The crux of our position is that CpG sites in genomes represent the places at which methylation can impact gene expression and thus selection; therefore, the number of CpG sites influences the ways by which gene expression can be influenced. As an analogy, we compare two stereo systems: the first has only one knob for volume, whereas the second has a knob for volume, but also has additional knobs for bass, treble, and balance. Although both stereos produce sound, the second system allows finer tuning, matching better the sound quality to the environment in which it is being played. CpG sites are similar to the knobs, which are adjusted via methylation; as the environment changes, knobs too are changed, increasing or decreasing gene expression (i.e., one aspect of sound quality). The elegance of epigenetic potential is that it requires no knob to be adjusted until the relevant environmental stimulus occurs; in other words, variation is latent until salient environmental cues release it. The more CpG sites a genome has, the more gene expression can be tuned and even reset to match the environment. In Kenyan house sparrows, we expect CpG sites to increase towards the range edge because dispersing individuals should benefit disproportionately from greater plasticity to cope with novel environments (30). Generally, though, we expect that epigenetic potential predominates in invasions as it allows for more and faster phenotypic change than through evolution alone, especially if bottlenecks and Allee effects occur. By contrast, as populations age and adapt, we expect that CpG site numbers will decline, fixing gene expression at one genetically accommodated level (31). In support in other systems, the abundance of CpG sites predicts levels of gene expression as well as high levels of variation in gene expression when organisms are exposed to different environments (32, 33).

Our epigenetic data are open to several interpretations, mostly because we were unable to discern what types of genomic regions were methylated, specifically how often methylation was associated with genes or gene regulatory regions. For instance, total DNA methylation decreased but variation in methylation increased towards the expanding range edge (Fig. 3*A*, *SI Appendix*, Fig. S3*A*). This result was unexpected but is consistent with other range expansions (34, 35). In those studies, low methylation in invading organisms was proposed to allow for the activation of genes necessary for invasion or facilitate the control of transposable elements, which could themselves increase genetic diversity and resultant phenotypic variation (34). We expect that until we can account better for context (i.e., gene, tissue, type of stimuli), it will be difficult to interlink epigenetic potential, methylation, gene expression, and phenotypic processes relevant to fitness (1). This unfortunately complex predicament is supported by our previous work in Kenyan house sparrows that found no detectable methylation signature across the range expansion, and modest differences between and native and introduced populations (18, 19). Other studies of other house sparrow introductions found similarly opaque results (36).

### The Loss of CpG sites and Genetic Assimilation

Another facet of our genetic data, however, implicate an important role for methylation in range expansion: its influence on mutation location and extent. Towards the expanding range edge in Kenya, we found that house sparrows lost few CpG sites to TpG and CpA mutations (Fig. 2*C* and 2*D*). Towards the core, by contrast, these mutations were comparatively common. These mutations, in particular their associations with CpGs and associated methylation, are suggestive of a mechanism of genetic assimilation. CpA and TpG mutations tend to be caused by persistent DNA methylation, such that CpG sites generally are four times less common than expected across the genome because 5-methylcytosines mutate four to fifteen times faster than any other genomic motifs (37–40). Historically, TpG and CpA mutations were thought to be completely random, but other mechanisms can promote deamination and mutation at methylated CpG sites (41–43). As a result, environmental conditions that induce DNA methylation at CpG sites can also lead to mutations (44, 45). To return to the stereo analogy, highly-methylated CpG sites might represent knobs that are tuned to a specific position and not changed for some time. Such knobs might eventually break off of the stereo, fixing that dimension of sound quality in place. In some cases, environmental extremes could expedite this fixation.

Further corroborating this perspective, we found that CpG sites proximal to sites that mutated in at least one individual or capture site exhibited the opposite trend seen for global DNA methylation: towards the range edge, methylation increased and variation in methylation decreased (Fig. 3*B*, *SI Appendix*, Fig. S3*B*). This pattern is intriguing because the methylation status of CpG sites can impact neighboring CpG sites (46). These effects may occur as unmethylated sites recruit demethylating enzymes and methylated sites recruiting methylating enzymes, which can act on proximal CpG sites (47). Towards the invasion core, we likely saw evidence of once consistently-methylated sites mutating and influencing the methylation status of neighboring CpG sites. As DNA methylation is likely to persist if the phenotype is advantageous, the more likely a specific CpG site is to be lost depends on deamination (48). Thus, high methylation at and near these sites could canalize gene expression. If methylation is extensive and enduring, over time what was once subject to plasticity might become genetically assimilated (48).

A final possibility involves the potentially critical nature of a minimum level of epigenetic potential for any colonization or related event associated with a small effective population size. Many natural and anthropogenic introductions fail, and whereas the causes of some failures are obvious, some are not (49). Epigenetic potential might represent a general mechanism whereby some small populations are able to surmount genetic bottlenecks, the accumulation of rare lethal recessive alleles, Allee effects or other phenomena associated with small founding populations (6, 50). Some threshold level of epigenetic potential might provide just enough latent phenotypic plasticity in contexts where increasing genetic diversity via recombination is improbable. In this light, epigenetic potential might be less of an individual trait promoting adaptation via plasticity than a mandatory condition for viability in a small or isolated population. We do not have the data to distinguish these alternatives here, but we expect that both will be important in the context of range expansions and comparable situations.

## Conclusions

Epigenetic potential represents the capacity for DNA methylation to occur within the genome and thus the range of phenotypic plasticity achievable. CpG sites can be inherited and selected for, allowing for epigenetic potential to increase if advantageous. As CpG sites are increased towards the range edge, we expect that they are advantageous to expanding individuals. Further, epigenetic mechanisms and phenotypic plasticity can act to preserve genetic variation, but persistent DNA methylation can contribute to CpG depletion (39, 51). The longer DNA methylation persists at a CpG site, the more likely that site is to deaminate and mutate (39). As DNA methylation is sensitive to environmental cues and can be dynamic throughout the life of an organism, mutation driven by epigenetic potential may be one mechanism by which environmental cues can alter the genomic architecture and lead to the assimilation of phenotype (48). The accumulation of mutations attributable to methylation towards the site of introduction suggests that this is occurring in across the Kenyan house sparrow range expansion. Altogether, epigenetic potential may represent a form of antifragility, a mechanism that buffers against unpredictable environmental conditions through phenotypic plasticity, eventually fixing genetic adaptations if conditions persist (52). We expect that epigenetic potential will be important in range expansion and invasion events broadly, and may be one reason house sparrows have achieved overwhelming success as an introduced species.

## Materials and Methods

### House sparrow sampling

House sparrows were captured via mist nets in Kenya in February through May of 2013. Individuals were brought into captivity and housed at ambient conditions with *ad libitum* access to food and water. After five days of captivity, house sparrows were euthanized via isoflurane overdose and rapid decapitation. Whole brains were removed and stored in PBS with sodium azide at −4°C. Before DNA extraction, hippocampi were excised from whole brain and 0.1 g was used for DNA extraction. Care was taken to extract the same region for each individual. DNA was extracted in August 2017 using phenol/chloroform/isoamyl and stored at −4°C until sequencing (53). Prior to sampling, all procedures were approved by the University of South Florida’s IACUC committee (W3877) and the Kenyan Ministry of Science and Technology.

### Sequencing

Sequencing was performed on an Ion Torrent Personal Genome Machine (PGM) at Georgia Southern University Armstrong Campus’ facility (Life Technologies). For library creation, we modified a standard GBS protocol for ddRAD and epiRAD sequencing on the Ion Torrent platform (Life Technologies) (25, 54). For ddRAD, we used enzymes *MspI* and *PstI* (all enzymes New England Biolabs, Ipswich, MA). For epiRAD we substituted the methylation sensitive restriction enzyme *HpaII* for *MspI*. After restriction digestion, we ligated on barcodes and y-adaptors of the Ion Torrent IonXpress sequences. We conducted emulsion PCR following manufacturers protocols of the Ion PGM-Hi-Q-View OT2-200 kit on the Ion Express OneTouch2 platform. We then sequenced resultant fragments following manufacturers protocols of the Ion PGM-Hi-Q-View Sequencing 200 Kit using an Ion 316v2 BC Chip. This process generated two datasets; the genetic ddRAD data that was used to determine epigenetic potential (*MspI* with *PstI*), and the epigenetic epiRAD data that was used to measure DNA methylation among HpaII restriction sites (*HpaII* and *PstI*).

### Genetic Data Quality Control and Analysis

The reads were demultiplexed within Torrent Suite™ version 4.4.3 (Life Technologies). The resulting BAM files were converted to SAM files using the “bamtosam” in the package *SAMtools* and imported into R (version 3.5.1) (55, 56). For each of the five sites, a density plot showing the distribution of read lengths against number of reads retained were created (*SI Appendix*, Fig. S6). The average read length peaked at 75, and all reads fewer than 75 base pairs were removed (*SI Appendix*, Fig. S6) (57). The resultant files were converted to FASTQ format and *TRIMMOMATIC* (version 0.36) was used with default settings to perform quality control (58). Reads were mapped back to the house sparrow genome using BWA (59, 60). The mapped reads were converted to a BAM file using *SAMtools* and run using STACKS pipeline, starting with the function “gstacks” to identify variant sites, followed by “populations” to calculate population genetic statistics (61). The Variant Call Format (VCF) file was returned from “populations”.

All CpG and GpC dinucleotides were identified within the mapped reads and the house sparrow genome. From CpG and GpC sites identified within the house sparrow genome, SNPs occurring within a CpG (e.g. CpG -> CpA) or within an GpC (e.g. GpC -> ApC) were considered a loss of that dinucleotide. Any SNPs leading to a distinct CpG or GpC motif to be formed were considered a gain of that dinucleotide. All mutation types leading to a loss or a gain of a CpG and a GpC were quantified as the average number of mutations: 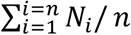 where *N*_*i*_ is the number of specific mutations within a sample, and *n* is the total number of samples. A Wilcoxon test was used to compare the significance of specific mutation types across all mutation types, and the FDR was calculated (*SI Appendix*, Fig. S2 and Table S2). In R, CpG sites were relativized using the equation:

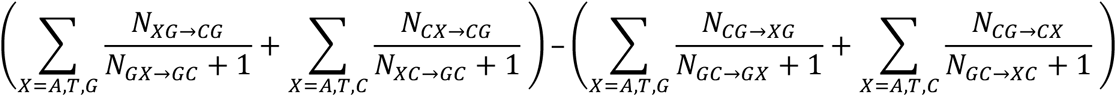

Losses were relativized using GpC losses:

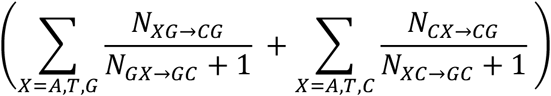

Gains were relativized using GpC gains:

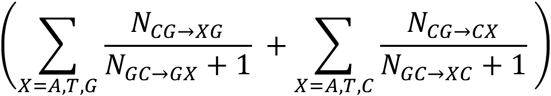

Number of CpA mutations were relativized using:

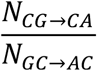

Number of TpG mutations were relativized using:

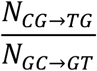

In all equations, X represents the changing base pair, N represents the number of mutations, and → represents mutation direction (28, 62).

Linear regressions were used to ask about the relationship between the distance from Mombasa as a fixed factor and relativized CpG sites, gains and losses of CpG sites, and CpA and TpG mutations.

A Discriminant Analysis of Principle Components was performed to elucidate population structure using data returned from *STACKS* (61, 63, 64). Within the package *adgenet*, the function “find.clusters” was used to predict the number of genetic clusters. The function “*dapc*” was then run to describe the relationship between clusters. Observed heterozygosity, expected heterozygosity, private allelic sites, and the inbreeding coefficient were compared to distance to Mombasa using Pearson correlation coefficients.

### DNA Methylation Quality Control and Analysis

Demultiplexed BAM files were returned from Torrent Suite™, converted to SAM format using SAMtools, and then to FASTQ format in the Linux environment (Torrent Suite) (55). The files underwent quality filtering using *TRIMMOMATIC* with default options (58). The files were mapped back to the house sparrow genome using *BWA* (59). BAM file was converted to BED file using the function “bamtobed” in *BEDTools* (65). BED files were merged into one feature used using “mergeBed” and the coverage of each loci was calculated for each house sparrow site using “coverageBed” in *BEDTools* (65). Any individual sparrows with fewer than 1000 reads were removed from the analysis (*SI Appendix*, Fig. S7). The median methylation level at each genomic location for all individuals was used to determine methylation value. A Wilcoxon test was used to detect loci that exhibited differential methylation between any two sampling sites, and the standard deviation was calculated for each individual bird. Within the loci that exhibited differential methylation, CpG sites within the same read segment as a loss or a gain of a CpG site were retained in a separate matrix. A Wilcoxon test was again used to test the difference in methylation levels at these specific loci between sampling sites, and the standard deviation was calculated.

## Acknowledgements

We thank the Martin lab for their many comments and suggestions, Kelly Hanson, Mark Jacim, and Jaime Zolik for help in the lab, and Alex Mutati for help with field collections. We would also like to thank the National Science Foundation (0920475, 1257773, 1656551, 1504662), and the USF Genomics Hub. A.L.L acknowledges support from the University of South Florida Graduate Student Research Challenge Grant. HEH acknowledges support from Sigma XI (G2016100191872782), the Porter Family Foundation, the American Ornithological Society Hesse Grant, and the American Museum of Natural History Frank M. Chapman Memorial Fund.

## Ethics Statement

All procedures were approved by USF IACUC (W3877 and IS0000636).

