## Supplementary material for "Epigenetic Potential and DNA Methylation in an Ongoing House Sparrow (*Passer domesticus*) Range Expansion": Fig. S1.

**This PDF file includes:**

Figures S1 to S7
Tables S1 to S2

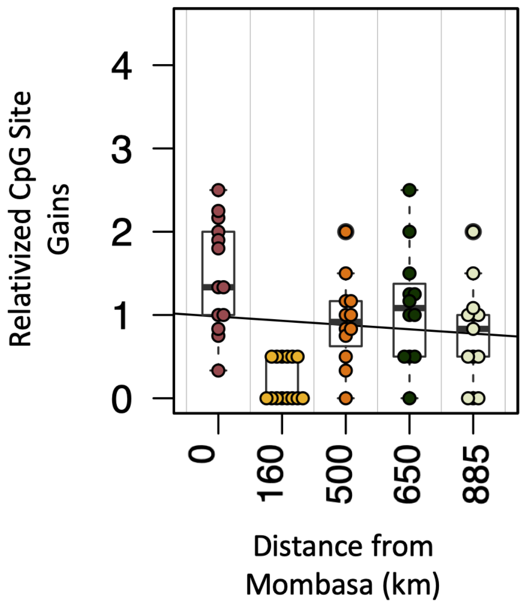


**Fig. S1. CpG site Gains.** There was no difference in gains of CpG sites (relativized using GpC gains) across the range expansion (R=-0.1, F-test p=0.41).

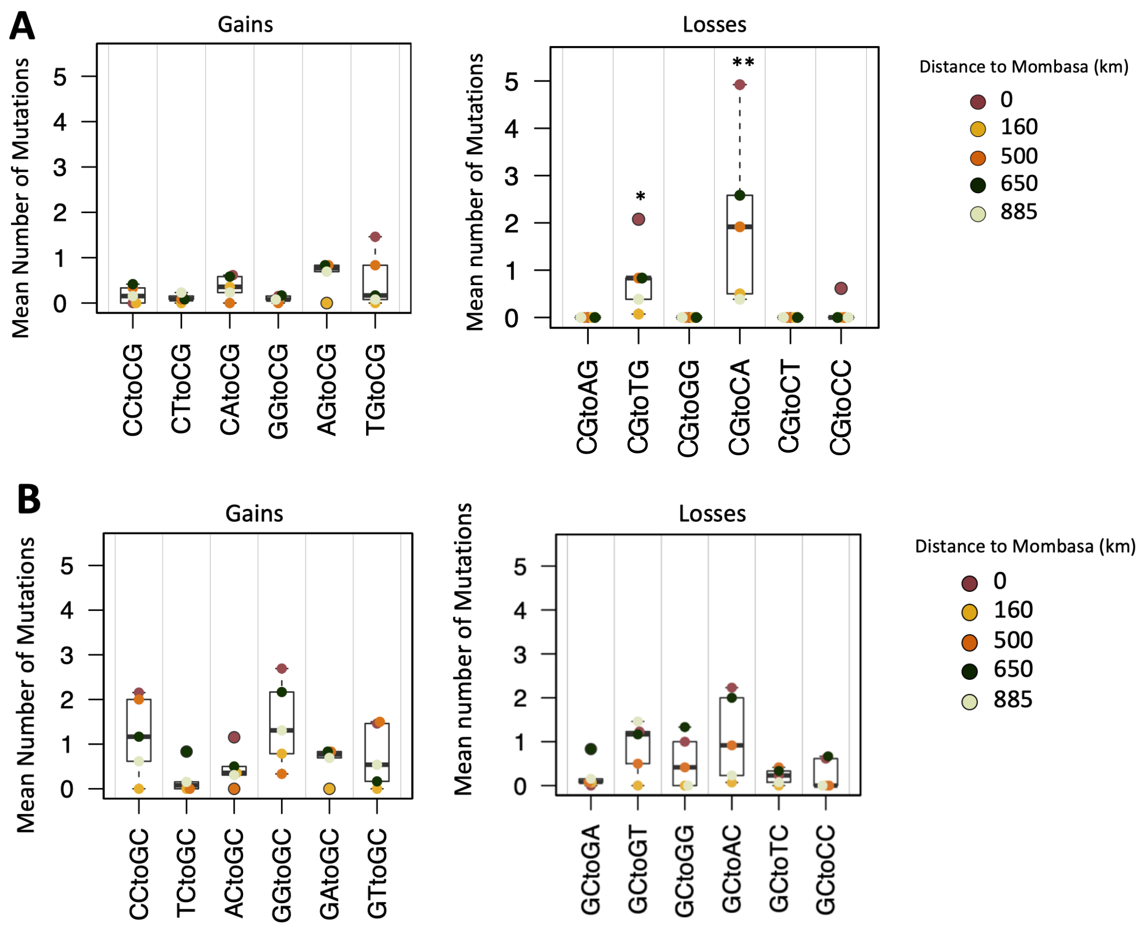


**Fig. S2. Average mutation types per sampling site.** Left panel represents mutations leading to a gain and right panel represents mutations leading to a loss of a CpG site (A) or GpC site (B). '*' represents FDR < 0.1, '**' represents FDR < 0.05.

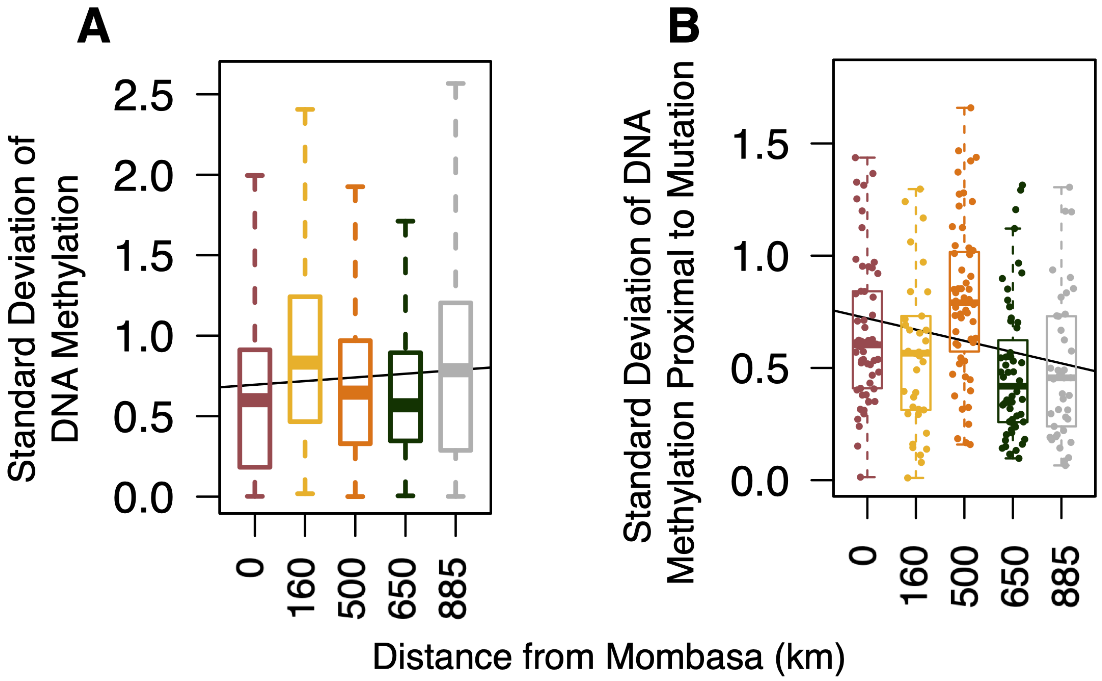


**Fig. S3. Variation in Methylation.** Standard deviation of DNA methylation calculated across the all CpG sites (A; R= 0.06, F test p= 3.008e-14) and at CpG sites proximal to mutated CpG sites (B; R= -0.21, F test p=0.003).

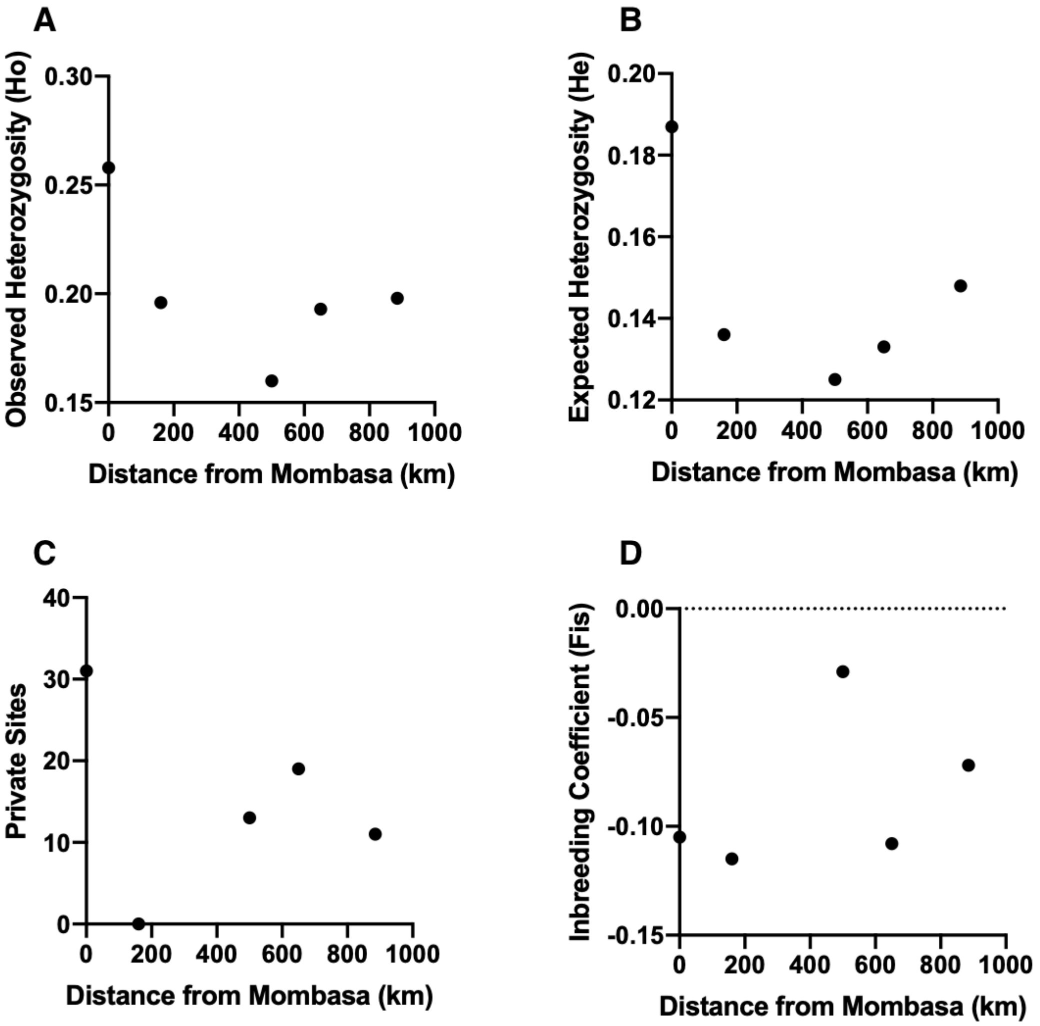


**Fig. S4. Estimates of genetic diversity compared to distance from Mombasa (km).** No estimate correlated with distance to Mombasa. (A) Observed heterozygosity (r= -0.5714, p= 0.3142), (B) Expected Heterozygosity (r= -0.5209, p= 0.3681); (C) Private allelic sites (r= -0.2385, p= 0.6993); (D) Inbreeding Coefficient (Fis) (r= 0.4174, p= 0.4844).

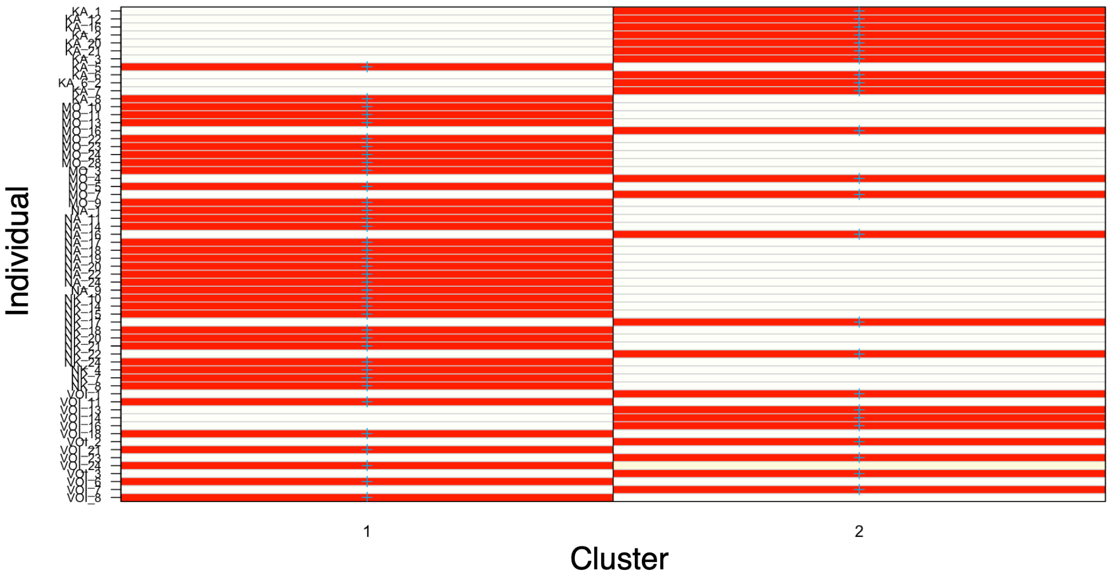


**Fig. S5. Discriminant analysis of principle components.** The assigned membership for each sample to one of the two predicted clusters. At least one individual from each sampling site was assigned to each cluster. In the individual IDs, sampling sites are abbreviated Mombasa (MO- 0 km), Voi (160 km), Nairobi (NA- 500 km), Nakuru (NK- 650 km), and Kakamega (KA- 885 km).

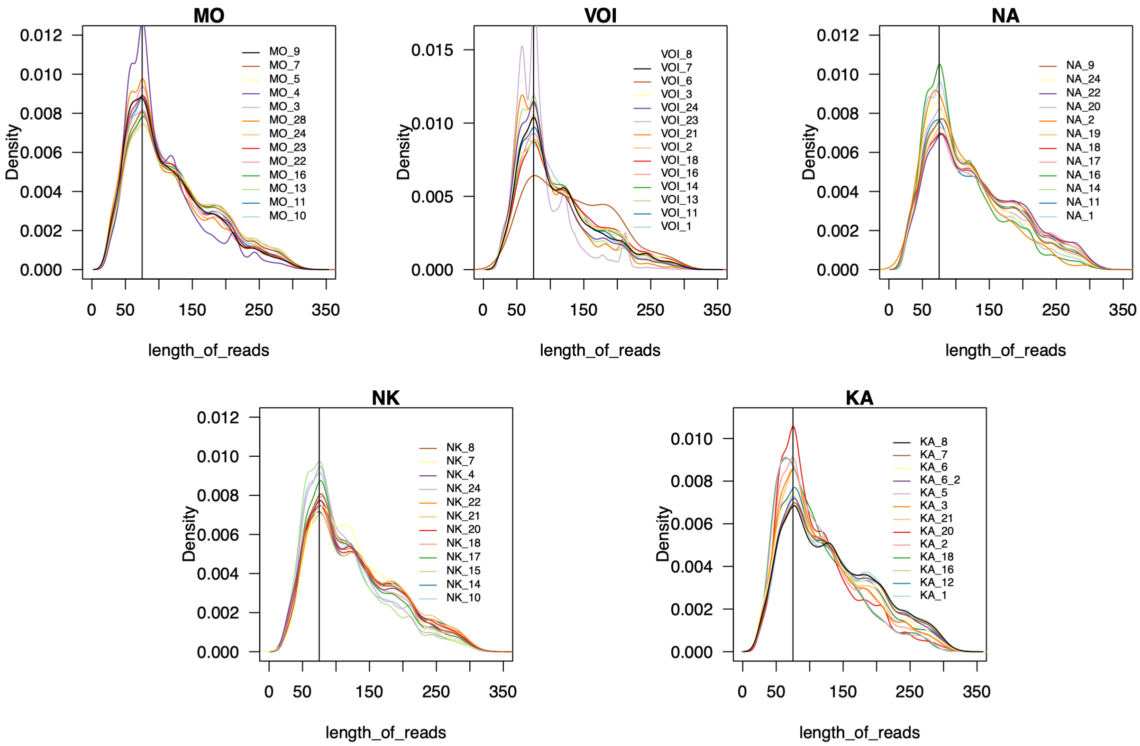


**Fig. S6. Density by read length plots.** Plot shows the distribution of RAD-Seq reads length in different populations. Sampling sites are abbreviated Mombasa (MO- 0 km), Voi (160 km), Nairobi (NA- 500 km), Nakuru (NK- 650 km), and Kakamega (KA- 885 km). The vertical line represents the read length equal 75bp.

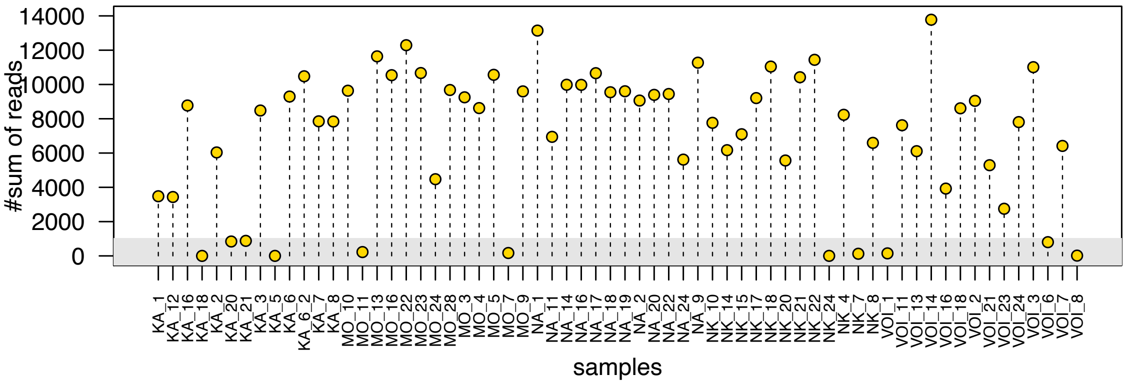


**Fig. S7. epiRADseq reads by individual.** Gray bar represents 1000 reads threshold. Samples with fewer than 1000 reads were not used in the epiRADseq analysis.

**Table S1. Sample sizes**

| Population | Distance from Mombasa (km) | Number of Individuals | |
|---|---|---|---|
| | | ddRADseq | epiRADseq |
| Mombasa | 0 | 13 | 11 |
| Voi | 160 | 14 | 11 |
| Nairobi | 500 | 12 | 12 |
| Nakuru | 650 | 12 | 10 |
| Kakamega | 885 | 13 | 9 |
| | **Total** | **64** | **53** |

**Table S2. Mutation type p-values and adjusted p-values**

| | Mutation Type | P-value | Adjusted p-value | | Mutation Type | P-value | Adjusted p-value |
|---|---|---|---|---|---|---|---|
| **Loss of CpG** | CG to AG | 0.991504646 | 0.991504646 | **Loss of GpC** | GC to GA | 0.906467106 | 0.946896667 |
| | CG to TG | **0.011870901** | **0.071225407** | | GC to GT | 0.173003242 | 0.519009727 |
| | CG to GG | 0.991504646 | 0.991504646 | | GC to GG | 0.631775109 | 0.946896667 |
| | CG to CA | **0.001018325** | **0.012219895** | | GC to AC | 0.115429686 | 0.461718744 |
| | CG to CT | 0.991504646 | 0.991504646 | | GC to TC | 0.904197667 | 0.946896667 |
| | CG to CC | 0.92474447 | 0.991504646 | | GC to CC | 0.946896667 | 0.946896667 |
| **Gain of CpG** | CC to CG | 0.588363185 | 0.946783864 | **Gain of GpC** | CC to GC | 0.093532894 | 0.461718744 |
| | CT to CG | 0.544456771 | 0.946783864 | | TC to GC | 0.942382574 | 0.946896667 |
| | CA to CG | 0.174837067 | 0.41960896 | | AC to GC | 0.656845893 | 0.946896667 |
| | GG to CG | 0.631189243 | 0.946783864 | | GG to GC | 0.015100796 | 0.181209556 |
| | AG to CG | 0.033732212 | 0.134928846 | | GA to GC | 0.368224891 | 0.736449782 |
| | TG to CG | 0.160829198 | 0.41960896 | | GT to GC | 0.333308784 | 0.736449782 |

Adjusted p-value is calculated using FDR.
